# Kinetics of osmotic stress regulates a cell fate switch of cell survival

**DOI:** 10.1101/2020.07.10.197871

**Authors:** Alexander Thiemicke, Gregor Neuert

## Abstract

Exposure of cells to diverse types of stressful environments differentially regulate cell fate. Although many types of stresses causing this differential regulation are known, it is unknown how changes over time of the same stressor regulate cell fate. Changes in extracellular osmolarity are critically involved in physiological and pathophysiological processes in several tissues. We observe that human cells survive gradual but not acute hyperosmotic stress. We find that stress, caspase, and apoptosis signaling do not activate during gradual stress in contrast to acute treatments. Contrary to the current paradigm, we see a substantial accumulation of proline in cells treated with gradual but not acute stresses. We show that proline can protect cells from hyperosmotic stress similar to the osmoprotection in plants and bacteria. Our studies found a cell fate switch that enables cells to survive gradually changing stress environments by preventing caspase activation and protect cells through proline accumulation.

## Introduction

All cells employ signal transduction pathways to respond to physiologically relevant changes in extracellular stressors, nutrient levels, hormones, and morphogens. These environments vary as functions of both concentration and time in healthy and diseased states ^1^. Cell signaling and cell fate responses to the environment are commonly studied using acute concentration changes ^1^. Only a few pioneering studies have explored the effects of the concentration and time, which is a gradual change of stimuli as a function of time on cell signaling in microbes ^2–5^ and in mammalian cells ^6–10^. Thus, the impact of the rate of environmental change on cell signaling, cell fate, and phenotype is a fundamental and poorly understood cell biological property (Fig. 1a). We address this lack in knowledge, by thoroughly measuring molecular changes in cells exposed to gradual environmental changes.

**Figure 1:**
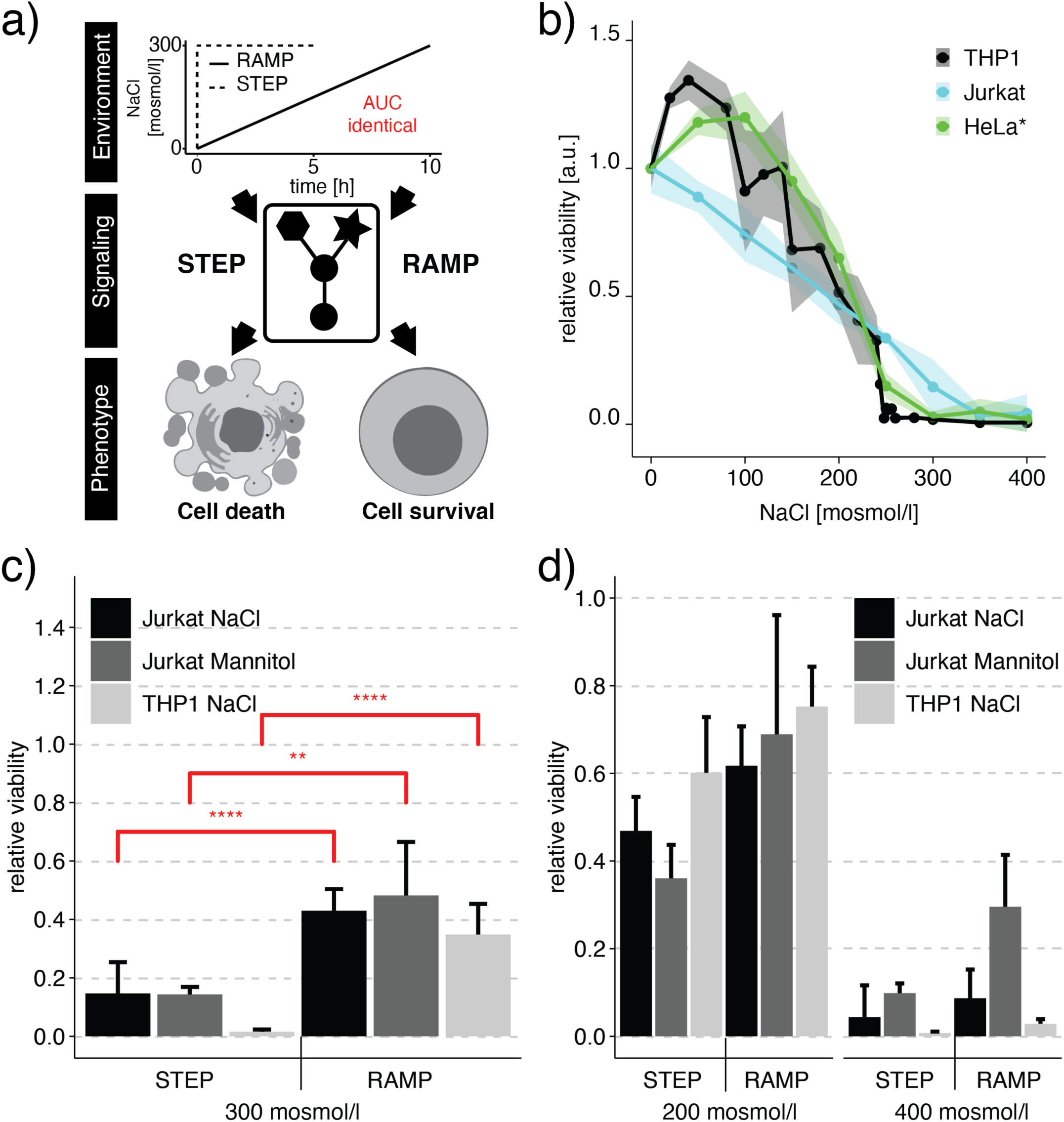
Human cell fate decisions are regulated differently upon step or ramp treatment conditions. a) Environments such as concentration ramps, as observed in different physiological relevant conditions, may differentially modulate cell signaling, cell fate, and phenotype even if the final concentration and total amount of stress are identical. Step experiments finish earlier than ramp experiments to account for the same total exposure or Area Under the Curve (AUC). b) We measured relative cell viability after exposure to instant hyperosmotic stress (NaCl for 5h for Jurkat, THP1) or 24h (HeLa cells). Cell viability was determined by measuring intracellular ATP (Jurkat, THP1) or cell counts (HeLa). The shaded area represents the standard deviation (SD) (Jurkat, THP1) or Standard Error (SE) (HeLa) (25) c,d) Relative cell viability was determined for step and 10h ramp treatment after addition of (c) 300 mosmol/l osmolyte or (d) 200 and 400 mosmol/l osmolyte. We determined viability at the end of the experiment after reaching the same cumulative exposure of additional NaCl. Bars represent data from at least 3 independent experiments for each condition. Error bars represent SD. Two-sided unpaired student’s t-test: **p<0.01, ***p<0.001, ****p<0.001.

To begin understand how the rate of environmental change regulates human cell fate decisions, we systematically expose cells to varying temporal profiles of increasing NaCl concentrations. NaCl is a ubiquitous osmolyte in the human body and causes cells to experience hypertonic stress at concentrations that change over time ^11–13^. While all tissues can experience increased NaCl concentrations in their microenvironment, measurements of osmolytes in the kidney have revealed very high physiological NaCl concentrations ^14,15^. In the kidney, spatial gradients of different osmolytes exist that change over time under normal and pathophysiological conditions ^16–18^. Hypertonicity changes over time are also known to occur in the intestinal system ^19,20^, the cerebrovascular discs ^21,22^, and the skin ^23^. In many of these high osmolarity tissues, resident immune cells provide basal protection or require migration upon an immune response of additional immune cells ^24^. Therefore, immune cells need to have the ability to survive such harsh high osmolarity environments that change over time. We choose immune cells as a model to systematically investigate how both rapidly and slowly increasing hypertonic, yet physiological environments impact cell survival, signaling, and metabolism.

## Results

### The rate of environmental change regulates cellular phenotype

We compared cell viability, cell signaling, and metabolism in cells exposed to either linear (ramp) or acute (step) concentration changes in the environments in which the final concentration and the total amount of osmotic stress (<u>A</u>rea <u>U</u>nder the <u>C</u>urve - AUC) is identical (Figure 1a). We identified the dynamic range of cell viability by determining the tolerance of monocytes (THP1 cell line, male, acute monocytic leukemia), T-cells (Jurkat, male, acute T cell leukemia), and cervical cells (HeLa, female, cervical adenocarcinoma ^25^) to step increases in NaCl concentrations (Figure 1b). In the non-stress control condition, cells were grown in culture under physiological NaCl concentrations of about 280 mosmol/l NaCl to which we added the hypertonic osmolytes NaCl and mannitol. To stress the cells and mimic *in vivo* osmolyte changes, we added up to 400 mosmol/l NaCl to the cells (Figure 1, Supplementary Figure 1). We observed that cell viability decreases with an increased NaCl concentration of up to 300 mosmol/l. At and above of 300 mosmol/l NaCl, cell viability is below 15% for all cell lines. Our results with the abovementioned cell lines are consistent with previous studies in HeLa cells (Figure 1b) ^25^, indicating that different cell types respond similarly to hypertonic stress.

We then quantified the response of different cell lines (Jurkat and THP1) to different rates and final NaCl concentrations (Figure 1c-d, Supplementary Figure 1). To compare the different conditions for the same final NaCl concentration, we exposed cells to the same cumulative exposure by integrating the total amount of NaCl over the entire profile (AUC). We performed experiments for each NaCl concentration for ramp durations of up to 10h. For experiments with ramp durations of less than 10h, cells stayed at the final NaCl concentration until the AUC is identical to the 10h ramp experiment. When we exposed Jurkat cells to 300 mosmol/l hypertonic osmolyte, viability improves from 15% to 40% for a ramp duration of at least 6h (Figure 1c (black), Supplementary Figure 1c (cyan)). In comparison, a step increase of 200 mosmol/l NaCl to the media for 5h reduced viability to around 50% and showed only minor improvement with increases in ramp duration (Figure 1d (black), Supplementary Figure 1c (magenta)). For the step condition of added 400 mosmol/l NaCl for 5 h, cell viability was below 5% and showed only minor improvement with increasing ramp durations (Figure 1d (black), Supplementary Figure 1 (green and yellow)). These observations are consistent in THP1 cells, indicating that this effect is reproducible in a different cell line and cell type (Figure 1c-d (light grey), Supplementary Figure 1a). To distinguish the effect on cell viability between NaCl toxicity and changes in external osmolarity, we repeated the experiments with mannitol in the Jurkat cell line at the same osmolar concentrations (Figure 1c-d (dark grey), Supplementary Figure 1b). Mannitol is not able to easily pass through the cell membrane and is known to have low cell toxicity. When we added 300 mosmol/l Mannitol to the medium, Jurkat cells survive better during the ramp compared to the step treatment. This comparison shows no difference between cells treated with NaCl or Mannitol, indicating extracellular hypertonicity and not NaCl-specific toxicity drive these effects. These results strongly suggest that cell viability improvements while slowly increasing NaCl concentration are a robust cell type- and cell line-independent hypertonic stress response.

### A functional temporal screen identifies regulators of cell viability in step and ramp conditions

Authors of previous studies argued that upregulation of genes encoding proteins responsible for the accumulation of cell internal osmolytes such as taurine (TauT), betaine (BGT1), sorbitol (AR) and inositol (SMIT) are the cause for improved viability in kidney cells exposed to a linear increase in osmolarity ^10^. To address if indeed these osmolytes are increased in our experiments, we determined the change in osmolyte levels in the cell by mass spectrometry measured in 5h step and 10h ramp conditions both to a final osmolarity of additional 300 mosmol/l NaCl. We found that sorbitol, inositol, betaine, taurine, and urea do not change compared to unstimulated cells (Figure 2a).

**Figure 2:**
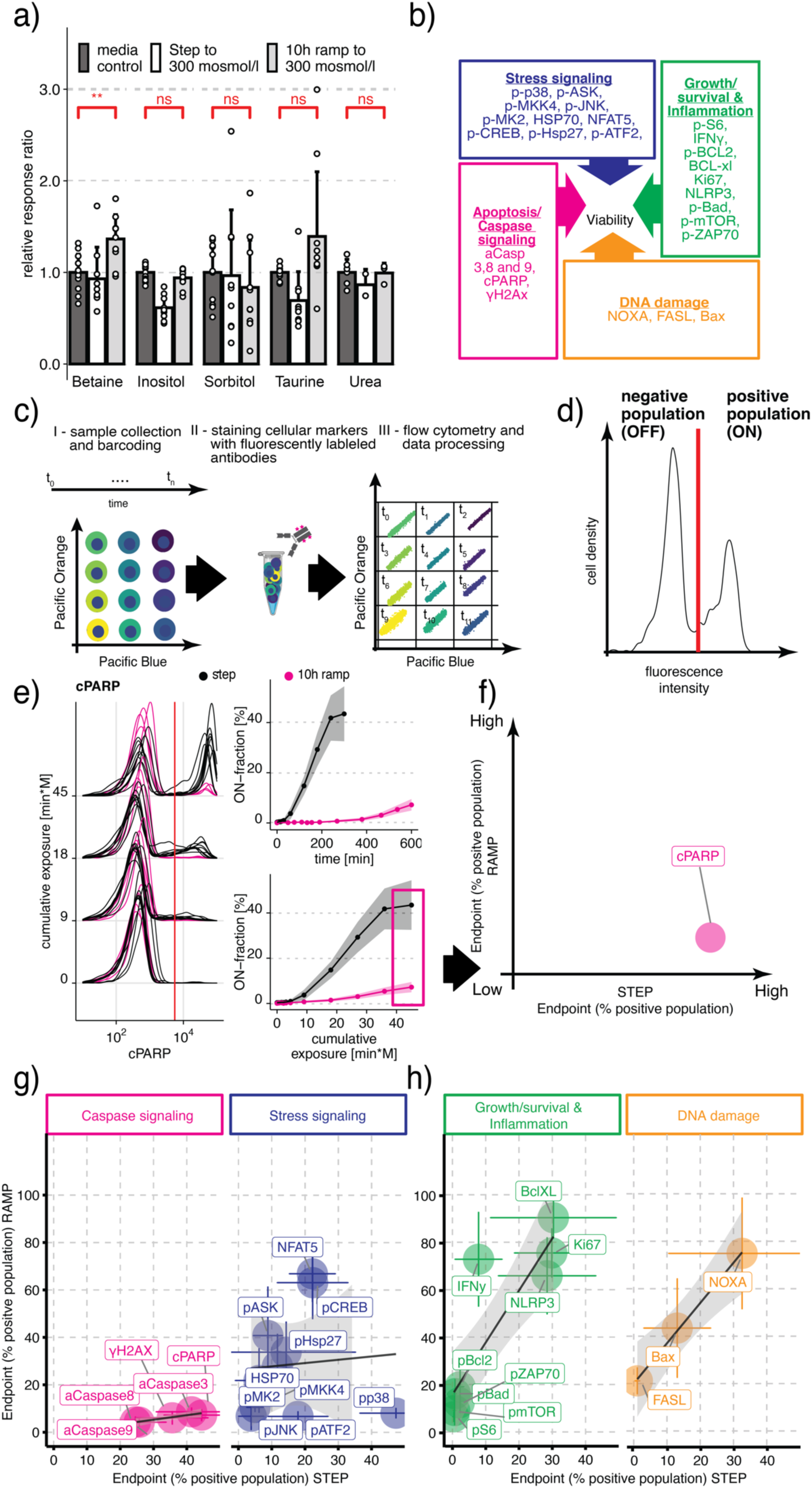
Temporal functional flow cytometry screen identifies differential regulation of stress and caspase signaling during step and ramp hyperosmotic stress conditions. a) Mean response ratio of cellular osmolytes relative to media measured in Jurkat cells exposed to an additional 300 mosmol/l NaCl and determined by Mass spectrometry. Two-sided unpaired student’s t-test: **p<0.01, ns=not significant. b) Overview of protein markers representing four cellular processes affecting viability. Each box lists the proteins representing each process. c) Multiplex flow cytometry workflow to quantify dynamic changes in protein activity over time: (I) Each time point is barcoded with a different combination of dyes. (II) Barcoded cells are pooled and split into different tubes for pairwise antibody staining. (III) We measured cells by flow cytometry and then computationally demultiplexed the different time points for further analysis. d) A single-cell distribution obtained by flow cytometry is threshold-gated (red line) to determine an ON-fraction. e) Representative flow cytometry single-cell distributions for cleaved PARP (cPARP) at selected time points for step (black) and 10h ramp (magenta) conditions (left). We quantified the fraction of cPARP positive cells (On-cells) as a function of time (right, top) or cumulative NaCl exposure (right, bottom). We plotted mean (solid line) and standard deviation (shaded area) of 3 – 10 biological replicas. f) We used endpoint measurement (magenta box in e) to determine ON-fraction to compare changes for step and ramp conditions. g,h) Comparison of endpoint measurement of mean ON-fraction between steps and ramps measured for individual markers of (g) caspase signaling (magenta), stress signaling (blue), and (h) DNA damage (orange), Growth/survival & Inflammation (green) in Jurkat cells in response to hypertonic stress. Circles represent the mean of 3-10 replicates per condition. ON-fraction at the final time point of cells exposed to 300 mosmol/l NaCl by a step (5h) or a 10h ramp (10h). Colored lines represent the SD. Black lines indicate linear regression fit lines. The shaded area represents 95% confidence interval.

To understand which cellular mechanisms contribute to improved viability during the slow ramp, we performed a temporal functional screen using a selected set of 27 well-established and validated markers of cell state and signaling that contribute to cell viability (Figure 2b). We grouped these into four cellular processes known to have an impact on cellular viability: stress signaling (blue), caspase signaling (magenta), DNA damage (orange) and growth/survival & inflammation (green) (Figure 2b). Each of these processes are known to be affected by increased NaCl concentrations ^11^. The process ‘stress signaling’ (blue) consists of markers belonging to stress/mitogen-activated protein kinases (SAPK/MAPK) pathways such as phosphorylated proteins p38 ^26,27^, JNK ^28^, MK2 ^29^, ASK1 ^30^, MKK4 ^31^, HSP27 ^32^, CREB ^33^, ATF2 ^34^, as well as protein levels of HSP70 ^35^ and NFAT5/TonEBP ^36,37^. MAPK pathways are known to convey stress signals to alter gene expression and cell phenotype ^38^. Proteins in the ‘caspase signaling’ group are initiator caspases^39^, such as activated caspase 8 (extrinsic pathway) ^40^ and caspase 9 (intrinsic pathway) ^40^, effector caspase 3^41,42^, cleaved PARP^42,43^ (cPARP) as a substrate of caspase 3 and histone H2AX (γH2AX), as a marker for the excessive DNA damage caused by DNA degradation during apoptosis^44^. The ‘growth/survival & inflammation’ group contain proteins that counteract apoptotic responses or indicate growth, proliferation, and inflammatory stimulation. The group contains phosphorylated forms of Bad^45^, Bcl2 ^46^, two pro-apoptotic proteins, mTOR ^47^, a key node in the cell growth pathway, ribosomal protein S6 ^48^, a marker for active translation, and p-ZAP70^49,50^, a marker for activated inflammatory signaling. The group also contains proteins Bcl-XL^51,52^, an anti-apoptotic protein, Ki67 ^53^, a general marker of a cell proliferative activity, and NLRP3 ^54,55^, a marker for the inflammasome, and intracellular IFNγ ^56^, a marker for inflammatory cytokine production. In response to DNA damage ^57^ proteins such as Noxa ^58–60^, Fas-L^60,61^, and BAX^59^ are expressed and fall into the group DNA damage.

We used fluorescent cell barcoding for multiplex flow cytometry to identify differentially regulated markers over time in step or ramp conditions ^62,63^. This functional temporal screen allows us to uniquely encode each time point sample with a combination of two dye concentrations (Figure 2c). We pooled barcoded samples and then split them again into different tubes to stain each split sample with specific antibodies. The advantages of first barcoding and then sample splitting are: reduced variability between samples; increased throughput; and reduced cost for different markers. Using this approach, we screened protein markers in Jurkat cells for their change over time in step versus 10h ramp experiments to an additional concentration of 300 mosmol/l NaCl. After data collection, we demultiplexed each sample with one or two protein markers to extract the individual time points (Figure 2c,d). To quantify each marker’s response, we next computed the fraction of positive cells for this marker and called this population ‘ON-fraction’ (Figure 2d,e). We then ploted the ON-fraction of each marker at the end of the time course experiment between the ramp and the step treatment to understand the correlation between the markers in each group (Figure 2f). This analysis revealed several distinct response patterns: (a) We observed strong activation in step but not ramp condition in cells with phosphorylated proteins of the caspase signaling group and p38 of the stress signaling group (Figure 2g, Supplementary Figure 2). (b) We observed minimal activation in step but strong activation in ramp conditions for some markers of stress response (pASK, NFAT5, and HSP70) (Figure 2g (blue), Supplementary Figure 3), growth (Ki67), anti-apoptotic (Bcl-XL), and inflammation (IFNγ, NLRP3) (Figure 2h (green), Supplementary Figure 4), and markers of DNA damage (Figure 2h (orange), Supplementary Figure 5). (c) A screen for other markers of cell survival, growth, and DNA damage reveals no significant differential changes over time. Based on this temporal functional screen, we focused on protein markers of the caspase signaling group.

### Caspases differentially regulate step and ramp conditions

Activated caspases 3, 8, 9, cleaved PARP and γH2AX all showed strong activation (ON-fraction) during the 300 mosmol/l NaCl step treatment (Figure 3a-e, black, Supplementary Figure 2e). Strikingly, caspase, and γH2AX activation, as well as PARP cleavage, are negligible during the 10h ramp treatment condition to the same final concentration (Figure 3a-e, magenta, Supplementary Figure 2e). Phosphorylation of γH2AX is also entirely prevented when caspase activity is inhibited during step NaCl treatment by a pan-caspase inhibitor (Supplementary Figure 6), which suggests prevention of apoptosis-associated destruction of DNA. Next, we investigated the contributions of caspases 3, 8, and 9 to the cell viability phenotype by quantifying the time course of activation for each member of the caspase signaling group relative to cleavage of PARP (Figure 3f). We found that caspase 3 (grey) is activated slightly before its target cPARP (purple), as expected (Figure 3f). Surprisingly, we found activation of the initiator caspases 8 (magenta) and 9 (cyan) after caspase 3 and cPARP. These results suggest that caspase 3 contributes to the induction of apoptosis, but not cleaved caspase 8 and 9. To understand if these population-level effects are indeed observable in the same cell, we co-stained cells with antibodies for activated caspase 9 and cPARP (Figure 4a). We found that single cells that are negative for cPARP are never positive for activated caspase 9 at any point during the treatment (Figure 4b). Cells positive for activated caspase 9 already have a high level of cPARP, suggesting that caspase 9 cleavage is not causative for apoptosis induction in single cells. Similarly, single cells co-stained for cPARP and activated caspase 8 are never negative for cPARP and positive for activated caspase 8, at the same time throughout the time course (Figure 4c). These results indicate no activation of caspase 8 before apoptosis induction (Figure 4d). In summary, these results suggest that activated caspase 3, but not activated caspase 8 and 9 contribute to PARP cleavage and subsequent induction of apoptosis (Figure 4d).

**Figure 3:**
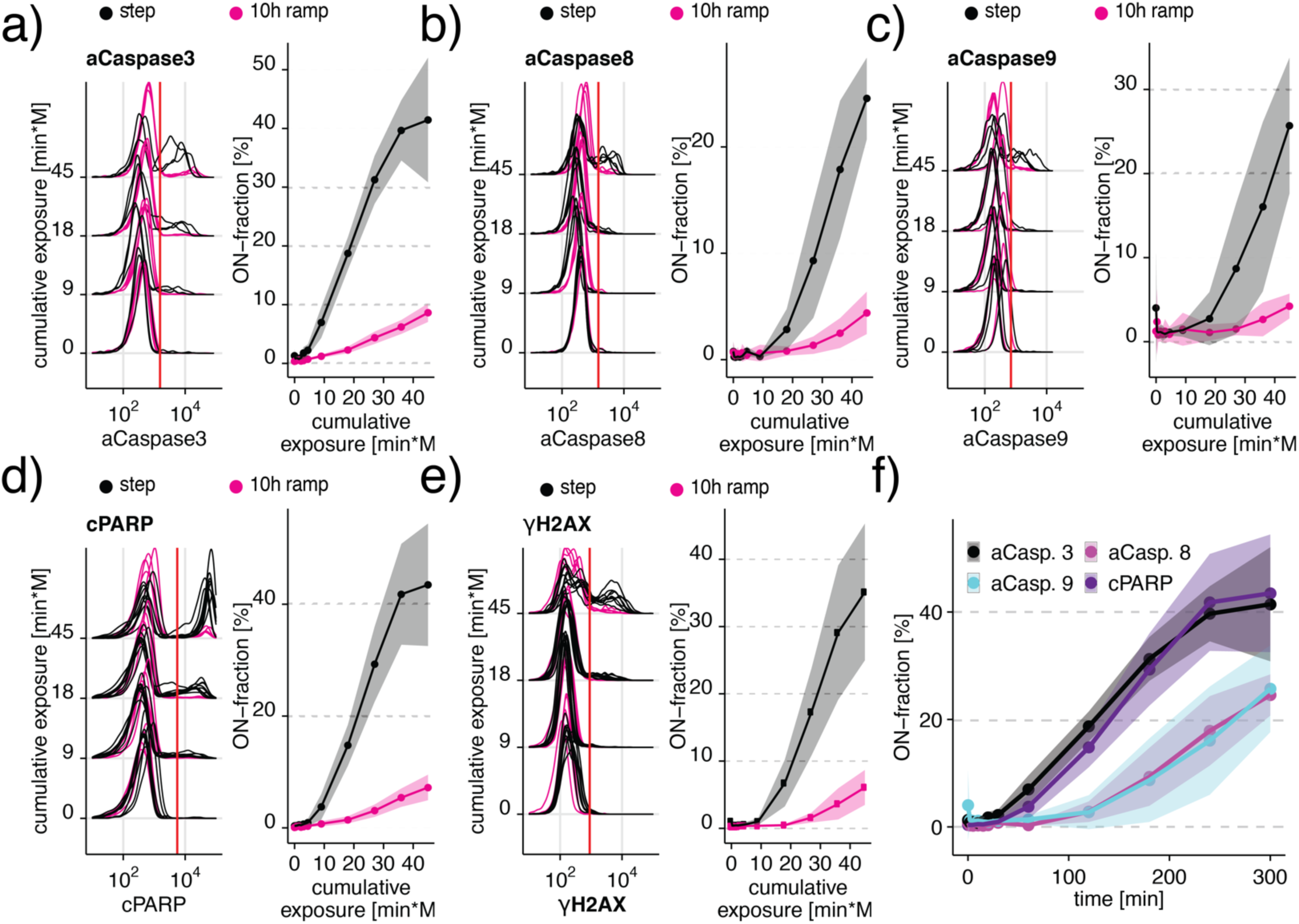
Differential caspase signaling regulates cell viability. a-d) Differential regulation of (a) cleaved Caspase 3, (b) cleaved Caspase 8, (c) cleaved Caspase 9, (d) cleaved PARP, and (e) γH2AX in Jurkat cells exposed to 300 mosmol/l NaCl by a step (black) or a 10h ramp (magenta). The left panel shows selected single-cell distributions over the cumulative exposure with individual lines representing independent experiments. Redline indicates the threshold for determining the ON-fraction. Right panels represent ON-fraction mean and standard deviation of 3-10 independent experiments as a function of cumulative exposure of NaCl. f) ON-fraction kinetics of caspase signaling markers over time indicate early (Caspase 3 and cPARP) and late (Caspase 8 and 9) activation. Lines indicate mean and SD of 3-10 independent experiments.

### Caspase signaling is the main contributor to cell death in step conditions

We next tested if these different caspases contributed to cell viability and addressed their mechanism in an attempt to link dynamics in caspase activation to apoptosis and cell phenotype (Figure 4e). In our ramp treatment condition to additional 300 mosmol/l NaCl in 10h, we found that cell viability increases to 40% in comparison to 15% in step treatment of the same final concentration and the total amount of NaCl relative to cells grown in control conditions (100% viability) (Figure 4e, magenta area). We asked if this increase in viability is entirely related to the lack of caspase activation and PARP cleavage, as observed in Figures 3 & 4. To test this idea, we treated cells with a step of 300 mosmol/l NaCl in the presence of different, potent pan-caspase inhibitors (panCas-i-a = Z-VAD-FMK^64^,panCas-i-b = Q-VD-OPH^65^) (Figure 4e). We observed an increase in cell viability to 40%, which is the same as for the ramp treatment. This result suggests that caspase activation and caspase-mediated apoptosis are necessary to explain the reduction in viability during the step treatment relative to the ramp treatment. Therefore we hypothesize that caspase-dependent apoptosis is the main contributor to the difference in viability between the step and the long ramp treatment conditions. We predicted that early caspase 3 activation triggers PARP cleavage and apoptosis compared to late caspase 8 and 9 activation (Figures 3, 4a-d). To test this prediction, we exposed cells to inhibitors of caspase 8, caspase 9 alone, or in combination. We found that inhibitors for caspase 8 and 9 do not substantially improve viability after step exposure to 300 mosmol/l NaCl (Figure 4e). As expected, we found that pan-caspase inhibition prevents the cleavage of caspase 3 during the step treatment (Figure 4f).

**Figure 4:**
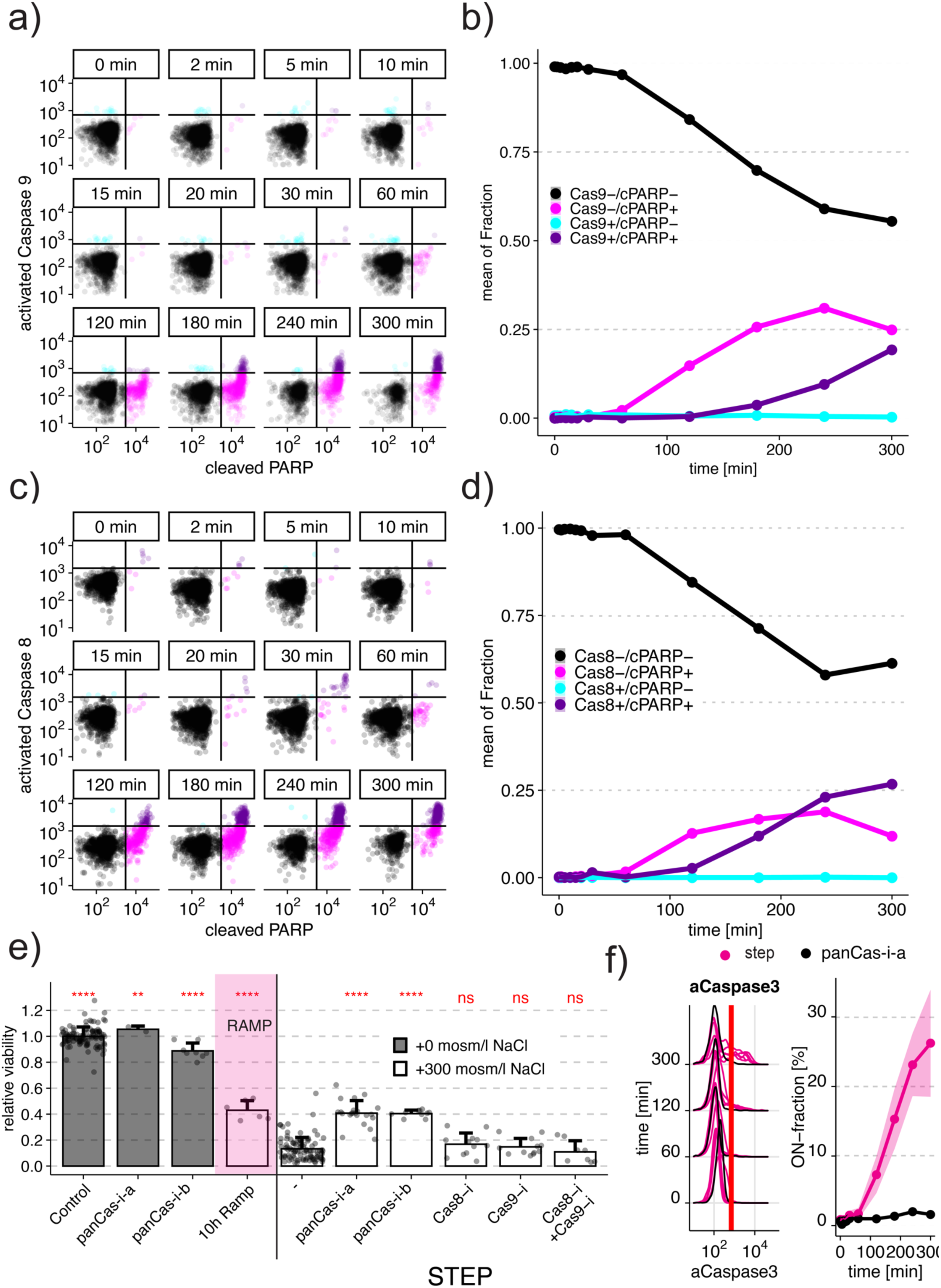
Activated Caspase 8 or 9 are not initiating apoptosis in hyperosmotic stress. a) Single-cell scatter plots of Jurkat cells co-stained with antibodies for cPARP and activated Caspase 9 measured by flow cytometry after exposure to 300 mosmol/l NaCl for 5h. Black lines indicate thresholds to determine individual fractions of low aCaspase9 and low cPARP (black), low aCaspase9, and high cPARP (magenta), high aCaspase9 and high cPARP (purple) and high aCaspase9 and low cPARP (cyan). Circles represent single cells. b) Quantification of fraction of cells stained for Caspase 9 activation and PARP cleavage over the time course using the thresholds indicated in (a). c) Single-cell scatter plots of Jurkat cells co-stained with antibodies for cPARP and activated Caspase 8 measured by flow cytometry after exposure to 300 mosmol/l NaCl for 5h. Black lines indicate thresholds to determine individual fractions of low aCaspase8 and low cPARP (black), low aCaspase8 and high cPARP (magenta), high aCaspase8 and high cPARP (purple) and high aCaspase8 and low cPARP (cyan). d) Quantification of fraction of cells stained for Caspase 8 activation and PARP cleavage over the time course using the thresholds indicated in (c). e) Relative viability of untreated cells (grey), cells exposed to a 10h ramp (magenta) or 5h step treatment both to 300 mosmol/l NaCl (white) exposed to different inhibitors. Inhibitors were added 30 min before NaCl at concentrations as follows: “panCas-i-a” (pan-caspase inhibitor Z-VAD-FMK, 100 μM), “panCas-i-b” (pan-caspase inhibitor Q-VD-OPH, 100 μM), “Cas8-i” (Caspase 8 inhibitor Z-IETD-FMK, 100 μM), “Cas9-i” (Caspase 9 inhibitor Z-LEHD-FMK, 100 μM). Bars indicate the mean and SD of at least 3 replicates. Two-sided unpaired student’s t-test: **p<0.01, ***p<0.001, ****p<0.001, ns=not significant. f) Activated Caspase 3 (aCasapse 3) in Jurkat cells exposed to 300 mosmol/l NaCl step in presence (black) or absence (magenta) of pan-caspase inhibitor (Z-VAD-FMK, 20 μM). The left panel shows single-cell distributions over the cumulative exposure with individual lines representing independent experiments. The Red line indicates the threshold for determining the ON-fraction. Right panels represent the mean and standard deviation of 1-4 independent experiments as a function of cumulative exposure.

Through our functional temporal screen, we also observed that p38 is strongly activated in NaCl step treatment condition, as previously reported to occur in other mammalian cells (Figure 5a) ^66,67^. However, during a 10h ramp treatment, we found that p38 is only slightly activated, perhaps playing a role in the decreased cell viability phenotype following step stimulation relative to the ramp stimulation. However, we found that the inhibition of all p38 protein isoforms by using a pan-p38 inhibitor (BIRB 796)^68^ had a statistically significant, but biologically small effect on cell viability following step treatment condition (Figure 5b). From these results, we conclude that the rate of hypertonic stress addition differentially regulates p38, but that p38 activity is not essential for the reduction in cell viability following step treatment.

**Figure 5:**
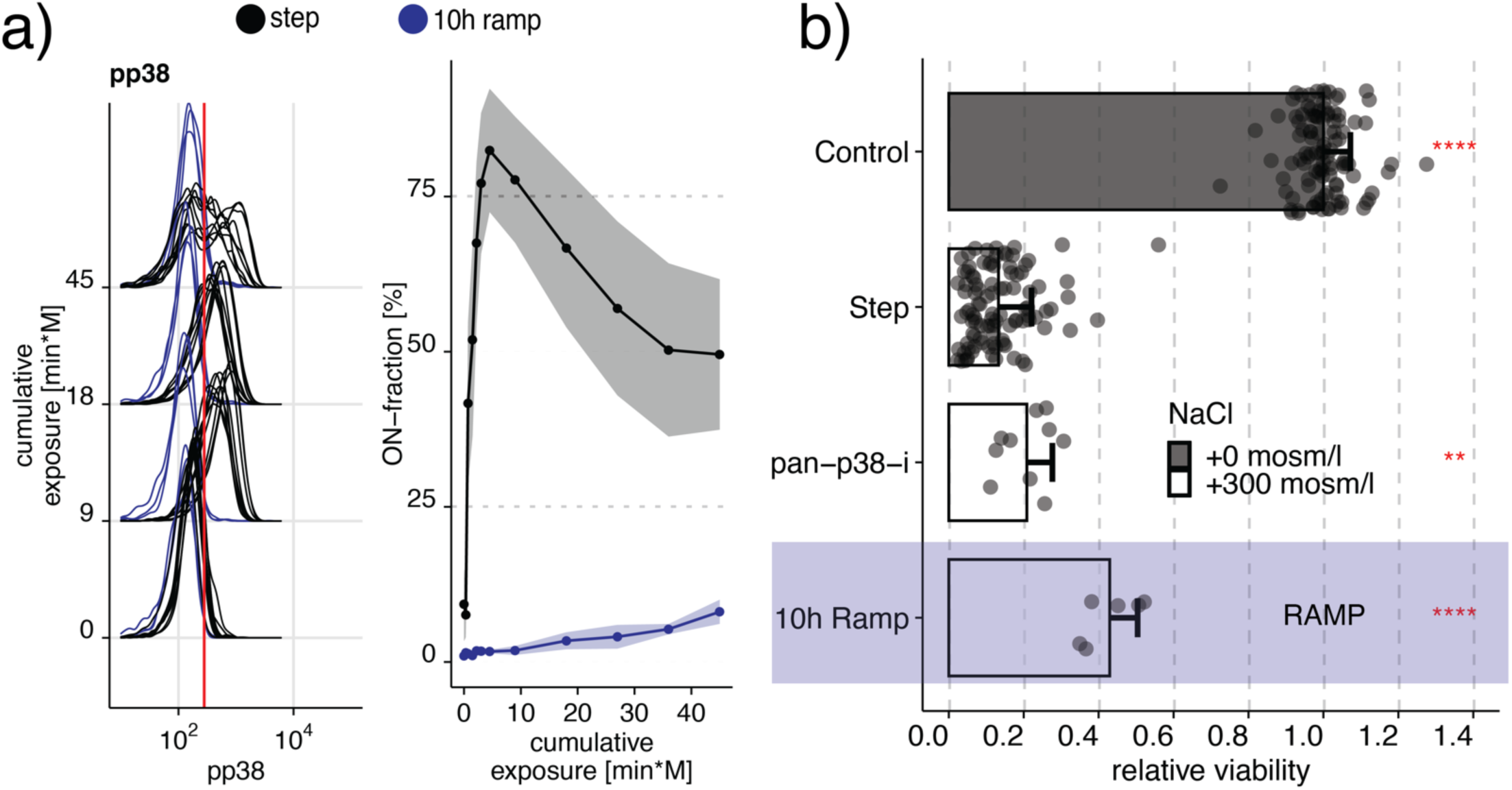
Contribution of p38 to apoptosis in hypertonic stress is minimal. a) Phosphorylation of p38 in Jurkat cells exposed to 300 mosmol/l NaCl by a step (black) or a 10h ramp (blue). The left panel shows selected single-cell distributions over the cumulative exposure with individual lines representing independent experiments. The Red line indicates the threshold for determining a cell that is p38 phosphorylation positive (ON-fraction). The right panel represents the ON-fraction mean and standard deviation of 3-10 independent experiments as a function of cumulative exposure. b) Viability of Jurkat cells relative to untreated cells (control) exposed to an additional 0 (grey) or 300 mosmol/l (white) NaCl for 5h (step) or 10h (ramp, purple), respectively. Pan p38 inhibitor (pan-p38-i, BIRB796) was added 30 min before NaCl at concentrations at 10 μM. Circles represent single experiments. Bars indicate the mean and SD of at least 3 replicates. Two-sided unpaired student’s t-test: **p<0.01, ****p<0.001.

Compared to p38, pASK, NFAT5, and HSP70 signals are reduced in step but not in ramp conditions shortly after osmotic stress (Supplementary Figure 3c,d,e). Followed by this initial drop are similar temporal profiles for step and ramp conditions. These results demonstrate that the dynamics of NFAT5, pASK, and HSP70 are not differentially regulated. We also observed similar dynamics for markers of the growth (Ki67), anti-apoptosis (Bcl-xL), inflammation (IFNγ, NLRP3), and the DNA damage (BAX, NOXA, and Fas-L) signaling groups (Supplementary Figure 4b, d, e, f). Markers that did change over time but not strongly between step and ramp conditions are the proliferation markers p-S6 and p-mTOR and pro-apoptotic protein p-BAD (Supplementary Figure 4a, g, h). We observed no change given the error in the measurements between step and ramp conditions for selected markers of stress signaling (p-MK2, p-JINK, p-MKK4, p-HSP27, p-ATF2, and p-CREB), and an pro-apoptotic protein p-Bcl2 (Supplementary Figures 3a, b, f, g, h, p, 4c).

### Intracellular proline levels improve viability in ramp stress conditions

To better understand the protective mechanisms contributing to improved viability during the ramp condition, we analyzed the abundance and fold changes of metabolites that may function as cell internal osmolytes (Figure 6a, Supplementary Figure 7). We found that among the most abundant metabolites are the amino acid proline, glutamic acid, and arginine. In comparison, traditional osmolytes such as betaine, inositol, sorbitol, taurine, or urea are significantly less abundant in the cell (Supplementary Figure 7). Interestingly, of these amino acids, only proline is differentially regulated in step and ramp conditions, rejecting the possibility that these amino acids are only byproducts of protein degradation (Figure 6a). This result suggests that proline may act as an osmoprotective molecule in human cells in ramp treatment conditions. The increase in abundance of cell internal proline levels relative to other amino acids and organic molecules suggests that cells import proline from the growth media. Elevated protein degradation in the cell, would presumably result in an equal distribution of increased amino acid abundance. We then tested if intracellular proline levels are independent of the activation of the caspase pathway or if preventing caspase-mediated cell death results in higher levels of proline in the cells. In these experiments, we exposed cells to a step treatment of NaCl with or without pan-caspase inhibitor Z-VAD-FMK (Figure 6b). As in all previous experiments, we exposed cells to the same cumulative exposure of NaCl for the same final NaCl concentration and compared the results. We found that regardless of pan-caspase inhibition, cells accumulated significantly less proline during the step treatment than cells exposed to the ramp treatment (Figure 6b). We conclude that caspase inhibition during hypertonic stress does not result in additional proline accumulation during the step treatment. This result indicates that caspase activation and proline accumulation are independent. To test if extracellular levels of proline can improve cell viability in the step treatment to 300 mosmol/l NaCl, we added free L-proline to the media of the cells before applying hypertonic stress (Figure 6c). We found a significant increase in viability due to added proline, in comparison to cells were no additional proline was added (Figure 6c). This result suggests that proline is transported into the cells and can protect mammalian cells from hypertonic stress. It is well established that hyperosmotic stress upregulates transporters for glutamine ^69,70^. Therefore, we tested if additional external L-glutamine, a precursor of proline^71^, can also improve viability. When we added additional L-glutamine to the media before adding NaCl, we observed a significant improvement in cell viability, similar to adding proline. Because proline is a yet unidentified compound in the mammalian response to hyperosmotic stress, we tested the effect of typical mammalian osmolytes on cell viability ^72,73^. When we added compounds identified as physiological osmoprotectants to the media, such as taurine, sorbitol, or betaine, we observe that these compounds seem to provide less or the same protection as proline or glutamine, during hypertonic stress. These results demonstrate that proline and glutamine are as effective as traditional osmolytes in protecting the cell from osmotic stress (Supplementary Figure 8).

**Figure 6:**
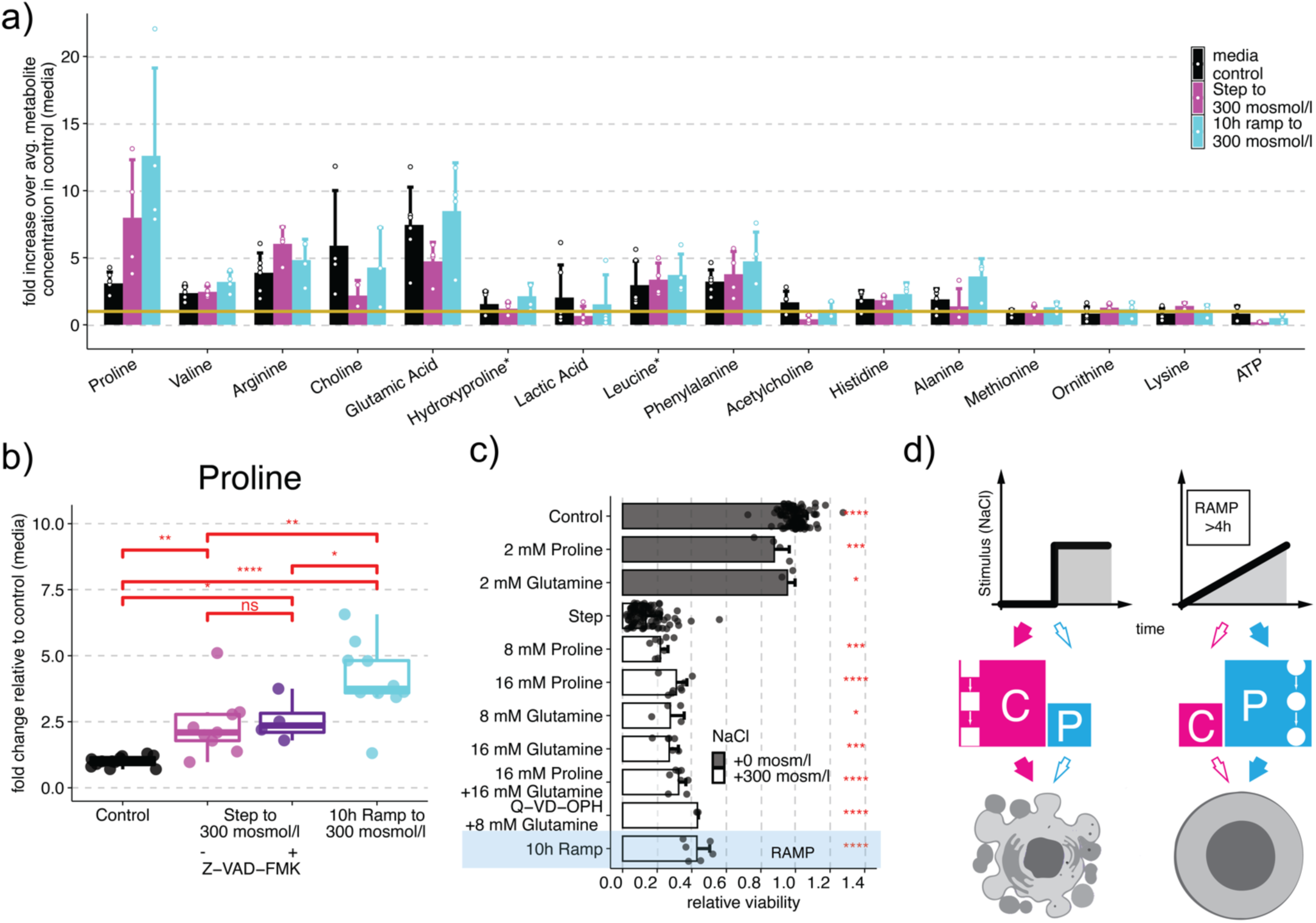
Intracellular proline protects human cells during ramp stress conditions. a) Fifteen most abundant metabolites detected in Jurkat cells without stimulation (control media, black), after treatment with step (magenta) or 10h ramp (cyan) to 300 mosmol/l NaCl. Bars represent the mean and standard deviation of the fold change of each metabolite to the average metabolite concentration in the control condition (yellow line) with circles representing individual replicates. b) Change of proline levels in Jurkat cells relative to control (no additional NaCl) in 0 (black) or 300 mosmol/l NaCl for 5h without (step, magenta) or with pan-caspase inhibitor “a” (Z-VAD-FMK, 100 μM)(purple) or a 300 mosmol/l NaCl ramp for 10h (cyan). Boxplots represent data of 4-10 replicates with circles represent individual replicates as determined by Mass spectrometry. c) External amino acid treatment impacts viability relative to untreated cells (control, grey) in Jurkat cells exposed to an additional 0 or 300 mosmol/l NaCl for 5h (step) or 10h (10h ramp, blue shade), respectively. Amino acids were added 60 min before NaCl at indicated concentrations. Pan-caspase inhibitor (Q-VD-OPH) was added 30 min before NaCl at 100 μM. Bars indicate the mean and SD of at least 3 replicates. Two-sided unpaired student’s t-test: *p<0.05, ***p<0.001, ****p<0.0001, ns=not significant. d) Model summarizing how instant stress conditions cause activation of caspase signaling (C) and cell death (left, magenta) whereas the gradual increase of the same stress to the same final concentration does not activate caspase signaling but instead increases intracellular proline (P) as an osmolyte to protect cells against increasing stress (right, cyan).

## Discussion

Previous studies have established that acute changes in environmental stimulus concentrations can control cell fate. However, cells in physiological environments may not necessarily experience such acute concentration changes. It is conceivable that typical solute concentration changes are gradual over time with different kinetics ^16,17,74,75^. However, there is a limited understanding of how a gradual change of stimulus concentrations affect cellular responses. We investigated stress responses of human immune cells to ramp increases in the concentrations of different osmolytes to address the key question of how varying the kinetics of stimulation affect cellular responses. We found that in comparison to instantly changing osmolyte concentrations, slow changes protect human immune cells from otherwise lethal insults (Figure 1b-d, Supplementary Figure 1). These results indicated that sensitivity to the rate of change of external osmolyte concentrations is a fundamental feature of human cells. These results are important because they demonstrate that immune cells that migrate into and through hypertonic tissues such as renal, intestinal, or epidermal tissue can survive hypertonic conditions better if these changes occur at a low rate over time. These results are consistent with pioneering studies indicating partial protection of renal medullary cells from slowly increasing external osmolytes ^10,14^. The authors of these pioneering studies postulate an increase in cell internal organic osmolytes is responsible for protecting cells exposed to gradually increasing osmolyte gradients ^10^. Perhaps surprisingly, we found that well-established osmolytes such as betaine, inositol, sorbitol, taurine, or urea did not increase at the end of our experiments (Figure 2a). One reason for this observation is likely that kidney cells respond differently to hypertonic stress than immune cells. Another reason is that we quantify cell internal osmolytes at the end of the 10h ramp experiment, whereas the previous study analyzed the response of cells 24h after the ramp treatment. We hypothesize that increases in traditional cell internal osmolytes after 24h may indeed function as a secondary and long-term protection against osmotic stress, but are not significant for short term protection. Because the step and ramp conditions do not differentially regulate the concentrations of these osmolytes (Figure 2), we studied the cellular pathways that are important in the regulation of cell viability during hyperosmotic stress. We discovered differential regulation between ramp and step conditions of caspases 3, 8, and 9 (Figure 3a-c). In step conditions a large fraction of cleaved caspases is observed, whereas in ramp conditions only a small fraction of cells show cleaved caspases (Figure 3a-c). This mechanism enables a population of cells to respond gradually to stresses that change over time without changing the ability of individual cells to undergo apoptosis. It is conceivable that in the kidney or the intestine, immune cells need to adjust not only to the absolute change but also to the rate of change in hypertonicity to avoid apoptosis. A property of an adapting system is to distinguish between a rapid and a slow increase of a stimulus. Adaptation has been studied in several important model systems, such as in yeast osmotic stress response signaling ^3–5,76^, chemotaxis signaling in bacteria ^2,77^, and mitogen^8,9,78^, and developmental^6^ signaling. These studies demonstrate that differential regulation of cell signaling between step and ramp stimulation might be a universal feature of signal transduction pathways by determining the presence or absence of a response to changes in the environment over time.

To better understand the mechanism of this observation, we analyzed the timing of caspase activation in single cells. We discovered that activated caspase 3 and cleaved PARP increase before activated caspase 8 and 9 (Figure 3f). These findings support previous studies demonstrating that activated caspase 3 cleaves PARP ^42,43^. This observation is consistent with published studies of apoptosis induction through caspase 9 protein recruitment, but not its cleavage. Recruited caspase 9 then cleaves caspase 3, which subsequently cleaves PARP ^39–41^. However, these cell population experiments cannot determine if indeed in a single cell, caspase 3 cleaves PARP and not caspase 8 or 9 (Figure 4a-d). To test if indeed caspase 3 cleaves PARP in single cells, we quantified co-stained cells for cleaved caspase 3 and cleaved PARP. Our single-cell analysis demonstrates that PARP gets cleaved before caspase 8 or 9, supporting our results and are consistent with previous cell population studies ^39,41^. From these single-cell time-course experiments, we predicted that inhibition of caspase signaling in step conditions increases cell viability similar to ramp conditions in single cells (Figure 4e). Because PARP activates before caspases 8 and 9, we predicted that these caspases do not contribute significantly to cell death. We indeed found that inhibiting caspase 8 or 9 individually, or together does not improve viability (Figure 4e).

To better understand which proteins contribute to differential caspase activation and cell survival, we analyzed changes in protein levels and/or phosphorylation states of upstream markers for proteins contributing to and indicating stress, growth, pro-apoptosis, anti-apoptosis, inflammation, and DNA damage. We separated these proteins into three groups. In the first group of protein markers of stress (NFAT5, pASK, and HSP70), growth (Ki67), anti-apoptosis (Bcl-xL), inflammation (IFNγ, NLRP3), and DNA damage (BAX, NOXA, and Fas-L) drop rapidly in step but not ramp conditions. These results could indicate that these markers can sense the difference in the type of stress gradient in a switch-like manner, although the dynamics of their distributions do not change overall. The second group of markers, such as proliferation markers p-S6 and p-mTOR and pro-apoptotic protein p-BAD, decreased over time but showed no differences between step and ramp conditions relative to the cumulative osmolyte exposure. These results indicate that a strong reduction of these markers is independent of the stress kinetics. The third group of proteins, such as stress signaling (p-MK2, p-JINK, p-MKK4, p-HSP27, p-ATF2, and p-CREB), and the pro-apoptotic protein p-Bcl2 did not show a clear difference between step and ramp treatments given the experimental constraints.

We also investigated the well-established link between osmotic stress and p38 signaling. We observed that p38 phosphorylation and phosphorylation of its target histone H2AX are also differentially regulated in ramp and step conditions (Figures 3e, 5). However, inhibition of p38 does not contribute to cell viability improvement as much as caspase inhibition (Figure 5b). These results are consistent with previous studies in macrophages where inhibition of stress response pathways such as p38 or JNK did not contribute to caspase signaling ^67^. This large temporal functional screen establish caspase signaling as the main contributor to differential regulation in step versus ramp stress condition compared to alternative signaling pathways of stress, proliferation, anti-apoptosis, pro-apoptosis, inflammation, and DNA damage.

Together these results indicate that human immune cells can survive shallow gradients to high osmolarity. This protective capability might be important because monocytes need to migrate inside the kidney from the low osmolarity cortex, to the very high osmolarity medulla to prevent bacterial infection ^81^. These results then beg the question of how do cells survive gradients of osmotic stresses that would otherwise be deadly?

We extended our initial analysis of cell internal organic osmolytes to a wide range of metabolites measured in step and ramp conditions. Although we detected many well-established osmolytes, their concentration is significantly lower than many other metabolites that we detected (Figure 2A, Supplementary Figure 7). Also, none of these osmolytes change significantly in step and ramp conditions (Figure 6). Instead, from this analysis, we discovered disproportional proline increases compared to the other amino acids. This disproportional increase for one amino acid excludes differential global protein degradation as a mechanism to increase proline levels (Figure 6a,b). Supplementing external proline or one of its precursors glutamine, protected cells from acute hypertonic stress, similar to stress protection in ramp conditions (Figure 6c). Although not well established in mammalian cells, in plants, proline acts as an osmoprotective molecule, and its accumulation is a well-described mechanism applied by plants to endure droughts and other stresses ^82,83^. Our results strongly suggest that the accumulation of intracellular proline plays a role in the protection of human immune cells from slowly increasing hypertonicity and the prevention of apoptosis (Figure 6c, Supplementary Figure 8).

In summary, we propose a model (Figure 6d) in which step increases in hypertonicity activate caspase signaling, PARP cleavage, and cause cell death. Whereas slowly increasing hypertonicity did not activate caspase signaling, but instead caused accumulation of intracellular proline. Proline is known to be upregulated during hypertonic stress in plants and bacteria to have an osmoprotective function. Proline functions as an organic osmolyte, molecular chaperone, metal chelator, and ROS scavenger independent of caspase activation ^82,84–86^. These properties make proline an efficient stress response molecule. We argue that proline has a underestimated and critical role in protecting human cells from cell death in hypertonic conditions and could explain how immune cells can survive in microenvironments within the body that have extreme osmolarities that change over time such as the renal papilla or the intestine.

## Methods

### Human cell culture

THP1 (ATCC® TIB-202(tm)) cells were cultured at 0.5–1 × 10^6 cells/ml in RPMI 1640 media (Corning, Catalog#: 15-040-CV) containing 10% Heat inactivated FBS (Gibco, Catalog#: 16140-071), 100 U/ml Penicillin-Streptomycin (Gibco, Catalog#: 15140-122), 2 mM L-alanyl-L-glutamine dipeptide (GlutaMAX™, Gibco, Catalog#: 35050-061) and 0.05 mM 2-Mercaptoethanol (Sigma, Catalog#: M3148) at 37 °C in a 5% CO2 humidity controlled environment. Jurkat cells (Clone E6-1, ATCC® TIB-152(tm)) and PBMCs (Stemcell technologies, Catalog # 70025.1) were cultured at 0.5–1.5 × 10^6 cells/ml in RPMI 1640 media (Corning, Catalog#: 15-040-CV) containing 10% Heat inactivated FBS (Gibco, Catalog#: 16140-071), 100 U/ml Penicillin-Streptomycin (Gibco, Catalog#: 15140-122) and 2 mM L-alanyl-L-glutamine dipeptide (GlutaMAX™, Gibco, Catalog#: 35050-061) at 37 °C in a 5% CO2 humidity controlled environment. Experiments with PMBCs were carried out 30 min after thawing.

### Experimental procedure for step and ramp stimuli application

A programmable pump (New Era Syringe Pump Systems, NE-1200) was used to apply gradually increasing (ramp) profiles. In brief, the pumping rate and dispensed volume per interval were calculated as described ^75^ and uploaded to the pump via a computer. A syringe pump driving a syringe (BD(tm), Catalog#: 309628) filled with 5 M NaCl (Corning, Catalog#: 46-032-CV) solution connected to a needle (Jensen Global, Catalog#: JG21-1.0x) with tubing (Scientific Commodities, Catalog#: BB31695-PE/4). The tubing was inserted into a foam stopper on an autoclaved glass flask (Pyrex, Catalog#: 4980-500) holding the suspension cells. Cells were shaken at 100 rpm during the entire experiment using a CO2 resistant shaker, ensuring proper mixing (Thermo Fisher Scientific, Catalog#: 88881101). For step stimulation, appropriate amount of 5 M NaCl (Corning® 500 mL 5M Sodium Chloride, #46-032-CV) solution was added by a syringe within 5 seconds to reach the desired final concentration. 5 ml of cells were removed with a syringe (BD(tm), Catalog#: 309628) through autoclaved silicone tubing (Thermo Scientific, Catalog#: 8600-0020) to collect time point samples.

### Cell viability assay

Cell viability was measured with CellTiterGlo (Promega, Cat.#: G7571). Cells were transferred to a white 96 well plate according to the manufacturer’s instructions and equilibrated to room temperature for 10 minutes. CellTiterGlo reagent was added in a ratio 1:8 to cell suspension. Luminescence was measured using a plate reader (Promega, GloMax Discover plate reader, GM3000). Relative viability was calculated by dividing luminescence values for each replicate by mean luminescence of media control for each experiment.

### Flow cytometry

Cells are fixed with 1.6% formaldehyde (Fisher, Catalog#: F79-4) in a 15 ml falcon tube. Fixation was quenched by adding 200 mM Glycine after 8 minutes. Cells were washed with PBS (Corning, Catalog#: 46-013-CM) and permeabilized with Methanol (Fisher, Catalog#: A454-4) for 15 minutes on ice. Cells were washed with PBS and stained with Pacific-Blue NHS ester (Pacific Blue(tm) Succinimidyl Ester, Thermo Fisher Scientific, #P10163) and Pacific-Orange NHS ester (Pacific Orange(tm) Succinimidyl Ester, Triethylammonium Salt, Thermo Fisher Scientific, #P30253) for 30 minutes. Cells are blocked with 1% BSA (Rpi, Catalog#: A30075-100.0) in PBS. Cells are washed and stained with a primary monoclonal antibody for 60 minutes at room temperature. Flow cytometry was performed on BD LSRII (five lasers). All antibodies used in this study are listed in supplementary Table 1.

### Flow cytometry analysis

Flow cytometry data was analyzed with custom R software. The primary cell population was gated on FSC-A vs. SSC-A by using the ‘flowcore’ package ^87^. The cell populations are automatically debarcoded and the resulting data was analyzed using custom software in R applying the following packages: ‘ggplot2’, ‘data.table’, ‘plyr’, ‘dplyr’, ‘flowViz’, ‘flowCore’, ‘flowStats’, ‘ggcyto’, ‘RcppEigen’, ‘fields’, ‘ggridges’, ‘viridis’, ‘scales’ and ‘xml2’. The distributions between independent experiments with similar shapes are aligned for their 0 minute time point so that their means are identical. This offset was applied to all the distributions in each experiment. Experiments are performed so that the total exposure to NaCl is identical between step and ramp experiments. The distributions, and ON-fraction are plotted as a function of the cumulative exposure. Plotting data as a function of the cumulative NaCl expose helps to distinguish between changes related to the total NaCl exposure compared to the temporal change in the NaCl concentration.

### Inhibitor studies

All inhibitors used in this study are listed in Supplementary Table 2. Inhibitors were dissolved in DMSO, and added 30 min before the start of the experiment to the cell culture media at indicated concentrations.

### Targeted Metabolomics Methodology

5 ml of cell suspension were pelleted, the supernatant was removed and resuspended in 90% methanol. Analysis of metabolites was performed at the Vanderbilt University Mass Spectrometry Research Center using an Acquity UPLC system (Waters, Milford, MA) interfaced with a TSQ Quantum triple-stage quadrupole mass spectrometer (Thermo Scientific, San Jose, CA), using heated electrospray ionization operating in multiple reaction monitoring (MRM) mode. 500 μl of each cell lysate sample was blown to dryness with N_2_ and reconstituted with 150 μL of an Acetonitrile/H_2_O (2:1) solution containing stable isotope-labeled internal standards: tyrosine-d_2_ and lactate-^13^C_3_ (Cambridge Isotope Lab, Tewksbury, MA). Centrifuged the cell lysate at 10,000 g for 20 minutes, and injected 90 μL supernatant into UPLC. The supernatant was chromatographically separated with a Zic-cHILIC column, 3 μm, 150 × 2.1 mm (Merck SeQuant, Darmstadt, Germany) at a flow rate of 300 μL/min. The mobile phases were A) 15 mM ammonium acetate with 0.2% acetic acid in water/acetonitrile (90:10, v/v), and B) 15 mM ammonium acetate with 0.2% acetic acid in acetonitrile/water/methanol (90:5:5, v/v). The gradient was as follows: 0 min, 85%B, 2 min, 85%B, 5 min, 30%B, 9 min, 30%B, 11 min, 85%B, 20 min, 85%B. We set the spray voltage to 5 kV and the capillary and vaporizer temperatures to 300°C and 185°C, with sheath gas and auxiliary gas set to 60 and 45 psi, respectively. The skimmer offset was −10 V, and the collision energy varied for each transition. Metabolites were identified based on predetermined peaks and elution times. The response ratio was calculated for each detected metabolite relative to the internal standard.

## Acknowledgments

This work is supported by NIH DP2 GM11484901 to GN, a predoctoral fellowship 18PRE34050016 from the AHA to AT, VICTR fund VR53716 to AT and Vanderbilt Startup Funds. The authors thank Robert Markowitz, Yelena Perevalova, and Minh Tran for technical assistance. The authors thank the Bachmann lab for technical assistance in setting up fluorescent cell barcoding for multiplex flow cytometry and the Vanderbilt Flow cytometry and Mass Spectrometry Core. The authors thank Dr. David G. Harrison, Dr. Jens Titze, Dr. Annet Kirabo, Dr. Meena S. Madhur, Dr. Vivian Gama and Dr. Jose Gomez for useful discussions and Benjamin Kesler, Jason Hughes, Dr. Hossein Jashnsaz, Dr. Amanda Johnson, Dr. Rama Ali, Dr. Rogert Colbran, Dr. David Cortez, Dr. Sandra S. Zinkel, and Dr. Ken Lau for feedback on the manuscript.

## Author information

Contributions

GN and AT conceived the study and designed the experiments. AT performed the experiments and the data analysis. GN and AT wrote the manuscript.

