## Supplementary_Information for "Kinetics of osmotic stress regulates a cell fate switch of cell survival"

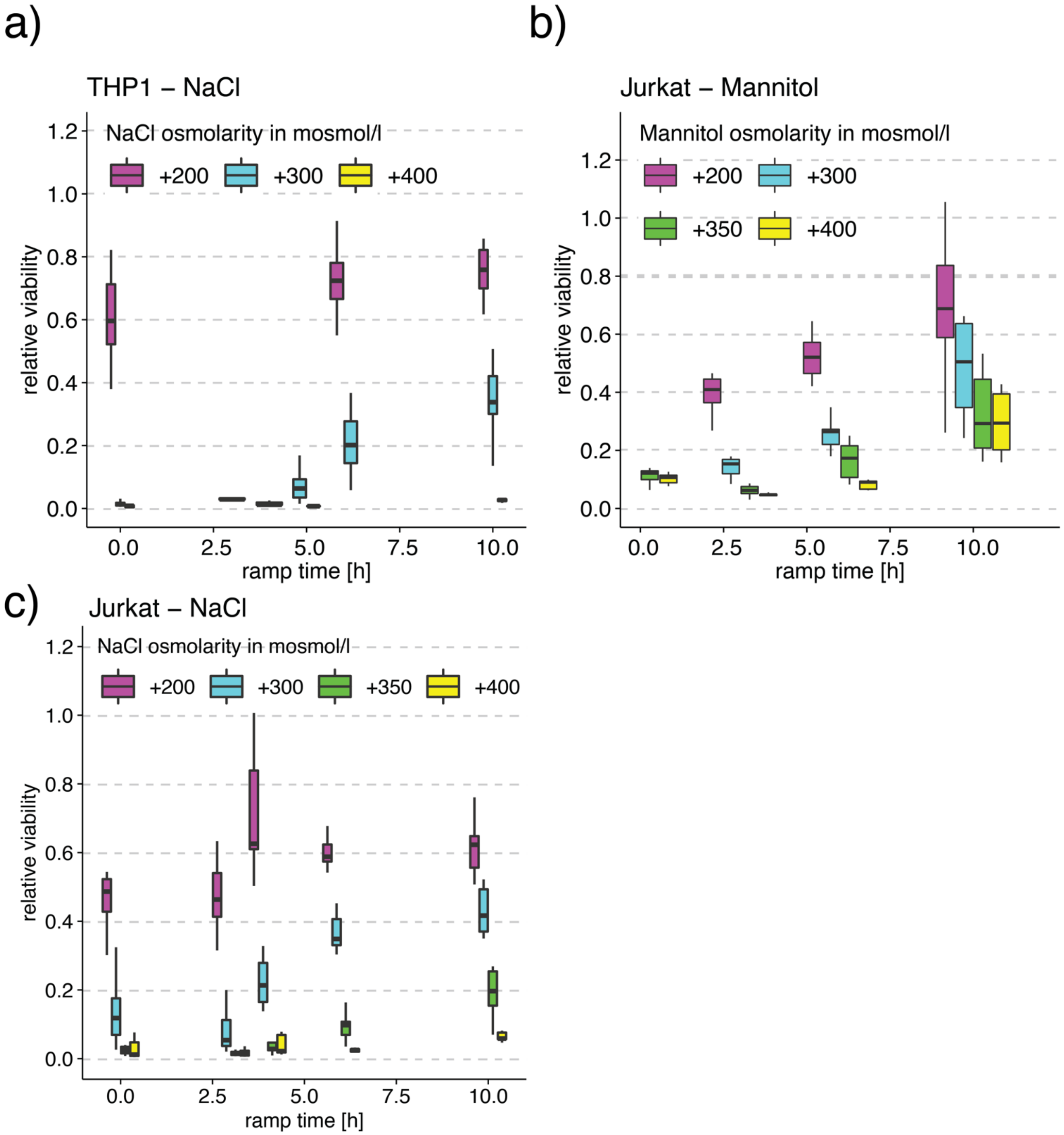


**Supplementary Figure 1: Viability improvement in hypertonic stress during a ramp vs. a step is a general cell biological feature independent of the cell line or osmolyte.** a) Viability for THP1 cells measured by intracellular ATP for step addition of NaCl (0), and ramps of 3,4,5,6,10h to indicated concentrations in mosmol/l. We determine viability at the end of the experiment at the same cumulative exposure of additional NaCl. Boxplots represent data from at least 3 independent experiments for each condition. b,c) Viability for Jurkat cells by measuring intracellular ATP for instant addition of mannitol (0), and ramps of 3,6,10h to indicated concentrations in mosmol/l for mannitol (b) and NaCl (c). We determine viability at the end of the experiment at the same cumulative exposure of additional mannitol. Boxplots represent data from at least 3 independent experiments for each condition.

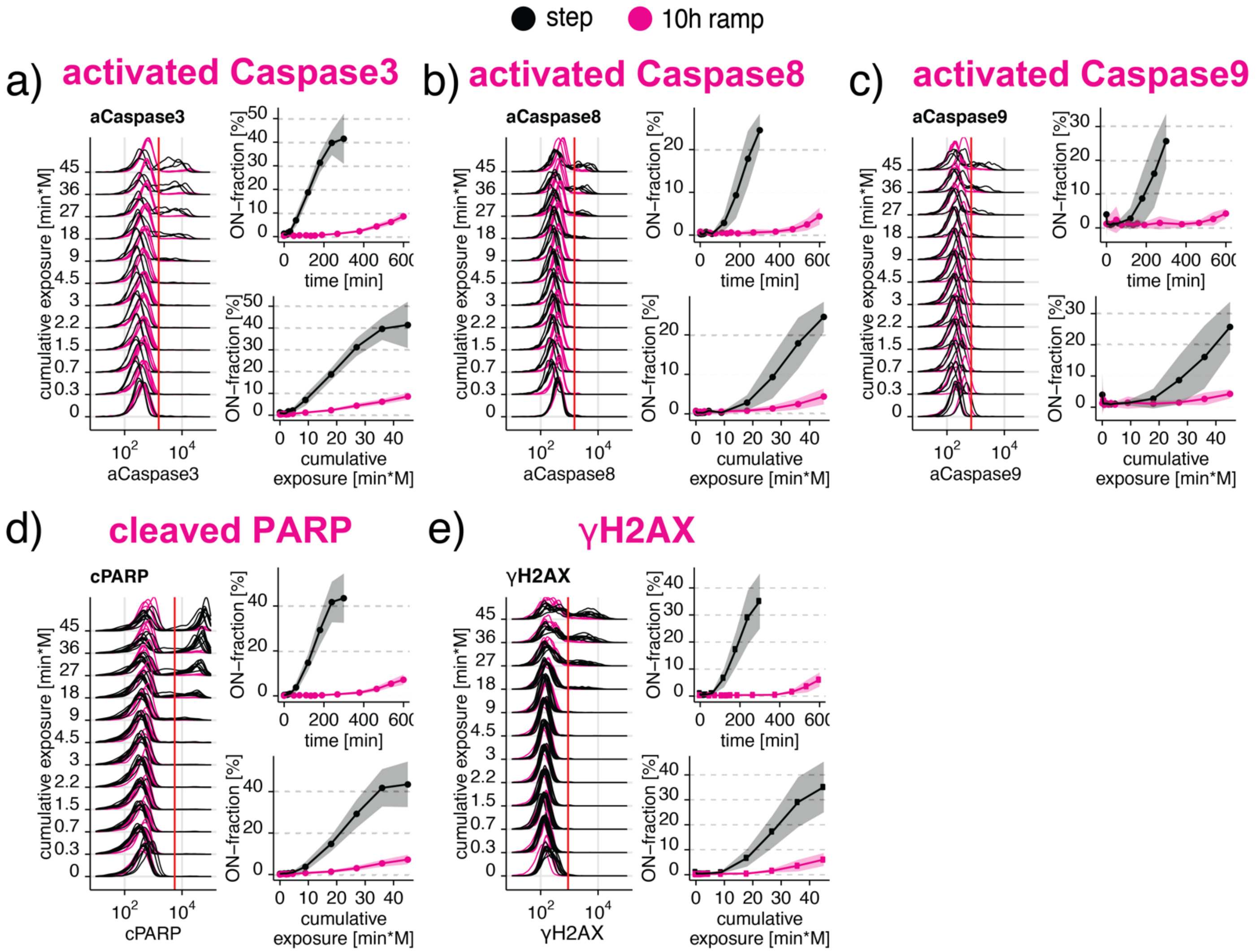


**Supplementary Figure 2: Differential caspase signaling regulates cell viability.** a-d) Differential regulation of (a) cleaved Caspase 3, (b) cleaved Caspase 8, (c) cleaved Caspase 9, (d) cleaved PARP, and (e) γH2AX in Jurkat cells exposed to 300 mosmol/l NaCl by a step (black) or a 10h ramp (magenta). The left panel shows single-cell distributions over the cumulative exposure with individual lines representing independent experiments. The Red line indicates the threshold for determining the ON-fraction. Right panels represent ON-fraction mean and standard deviation of 3-10 independent experiments as a function of time (top panel) or cumulative exposure of NaCl (lower panel).

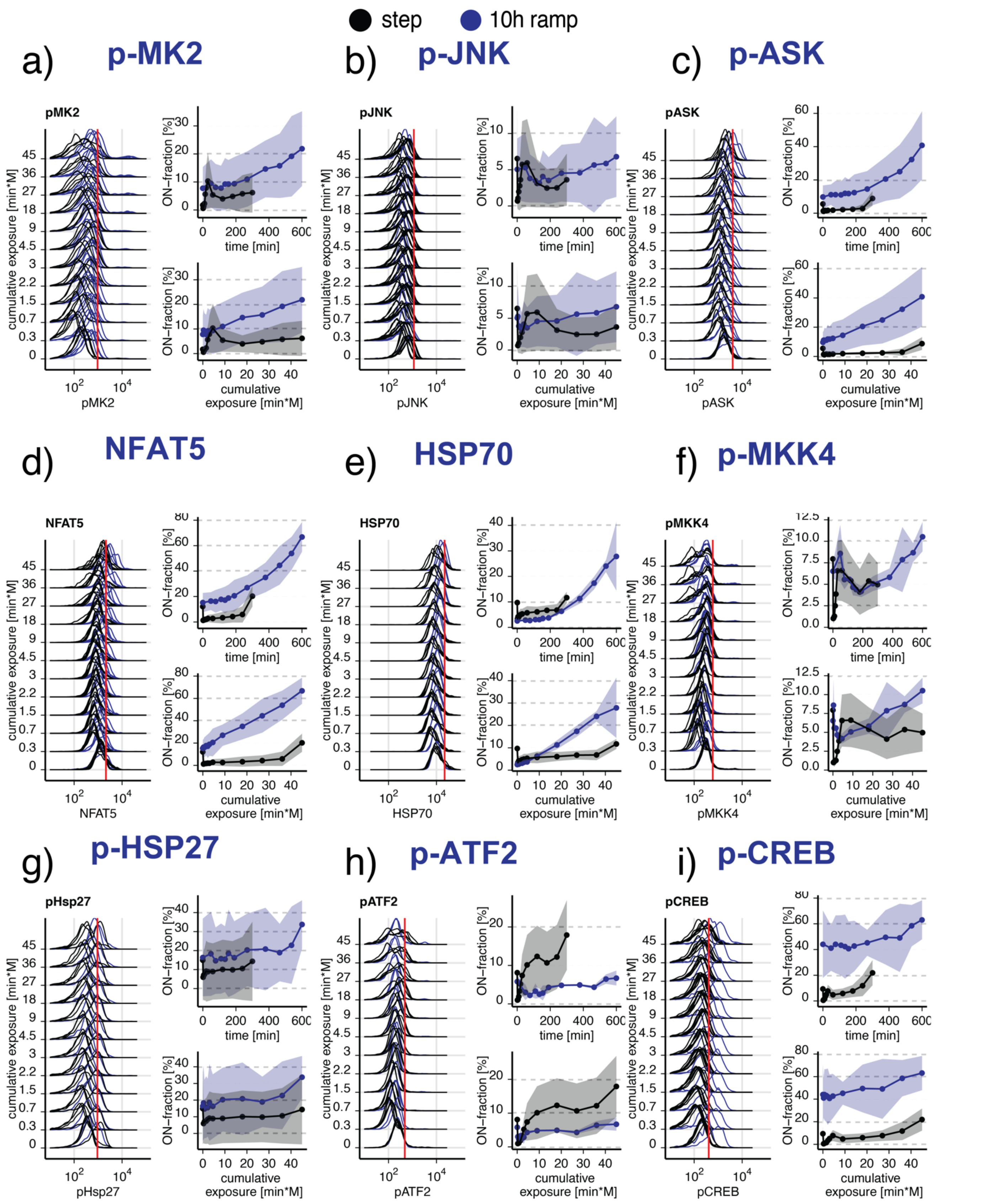


**Supplementary Figure 3: Markers for stress signaling.** a-i) Markers for stress signaling in Jurkat cells exposed to 300 mosmol/l NaCl by a step (black) or a 10h ramp (blue). The left panel shows single-cell distributions over the cumulative exposure with individual lines representing independent experiments. The Red line indicates the threshold for determining the ON-fraction. Right panels represent the mean and standard deviation of 3-10 independent experiments over time (top panel) and cumulative exposure (lower panel).

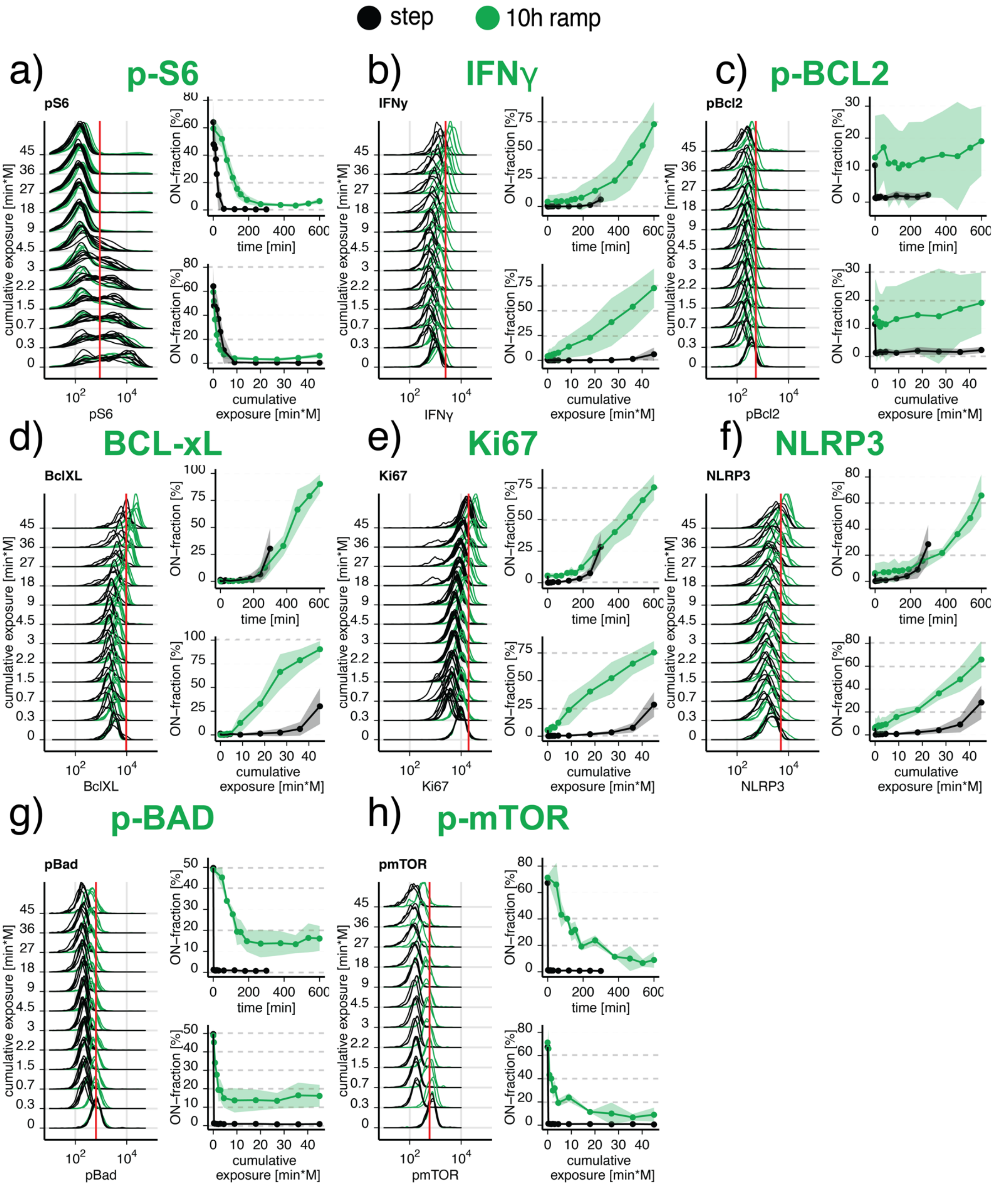


**Supplementary Figure 4: Markers for growth and proliferation.** a-h) Markers for growth/proliferation in Jurkat cells exposed to 300 mosmol/l NaCl by a step (black) or a 10h ramp (green). The left panel shows single-cell distributions over the cumulative exposure with individual lines representing independent experiments. The Red line indicates the threshold for determining the ON-fraction. Right panels represent the mean and standard deviation of 3-10 independent experiments over time (top panel) and cumulative exposure (lower panel).

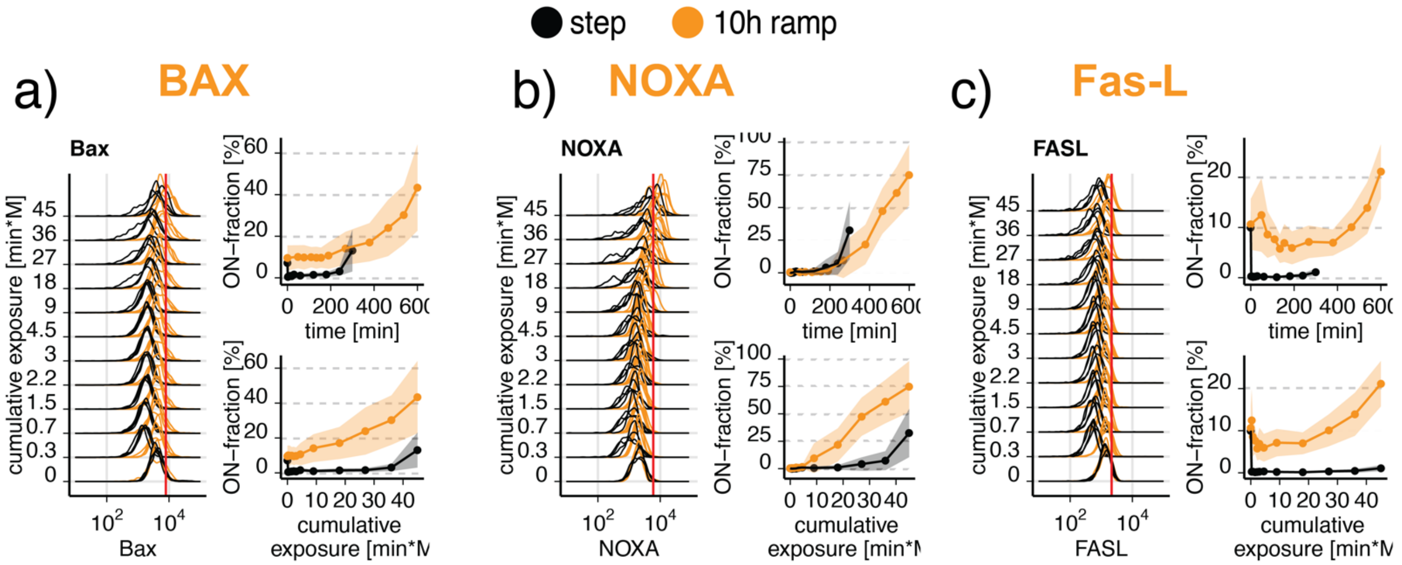


**Supplementary Figure 5: Markers for DNA damage.** In a-e) Markers for DNA damage in Jurkat cells exposed to 300 mosmol/l NaCl by a step (black) or a 10h ramp (yellow). The left panel shows single-cell distributions over the cumulative exposure with individual lines representing independent experiments. The Red line indicates the threshold for determining the ON-fraction. Right panels represent the mean and standard deviation of 3-10 independent experiments over time (top panel) and cumulative exposure (lower panel).

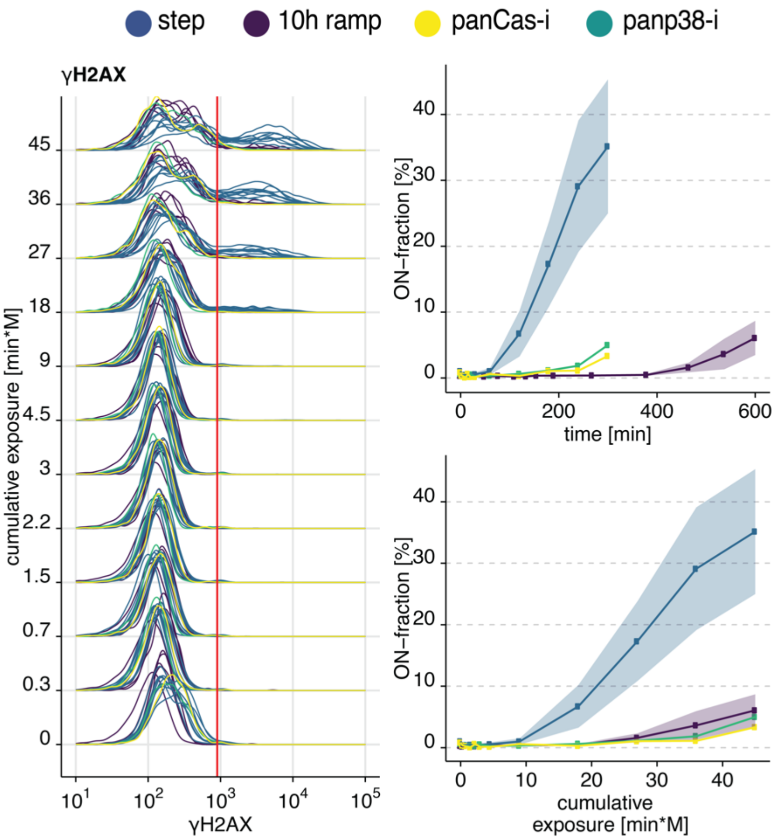


**Supplementary Figure 6: Caspases and p38 mediate γH2AX phosphorylation during hyperosmotic stress.** γH2AX phosphorylation in Jurkat cells exposed to 300 mosmol/l NaCl as a step without (blue), with inhibitor pan-Cas-I (Z-VAD-FMK, yellow), with inhibitor pan-p38-I (BIRB796, green), or as a ramp for 10h (purple). Left panel shows single-cell distributions over the cumulative exposure with individual lines representing independent experiments. The Red line indicates the threshold for determining the ON-fraction. Right panels represent the mean and standard deviation of 1-10 independent experiments over time (top panel) and cumulative exposure (lower panel). Z-VAD-FMK (panCas-i) was added 30 min before NaCl at 20 μM.

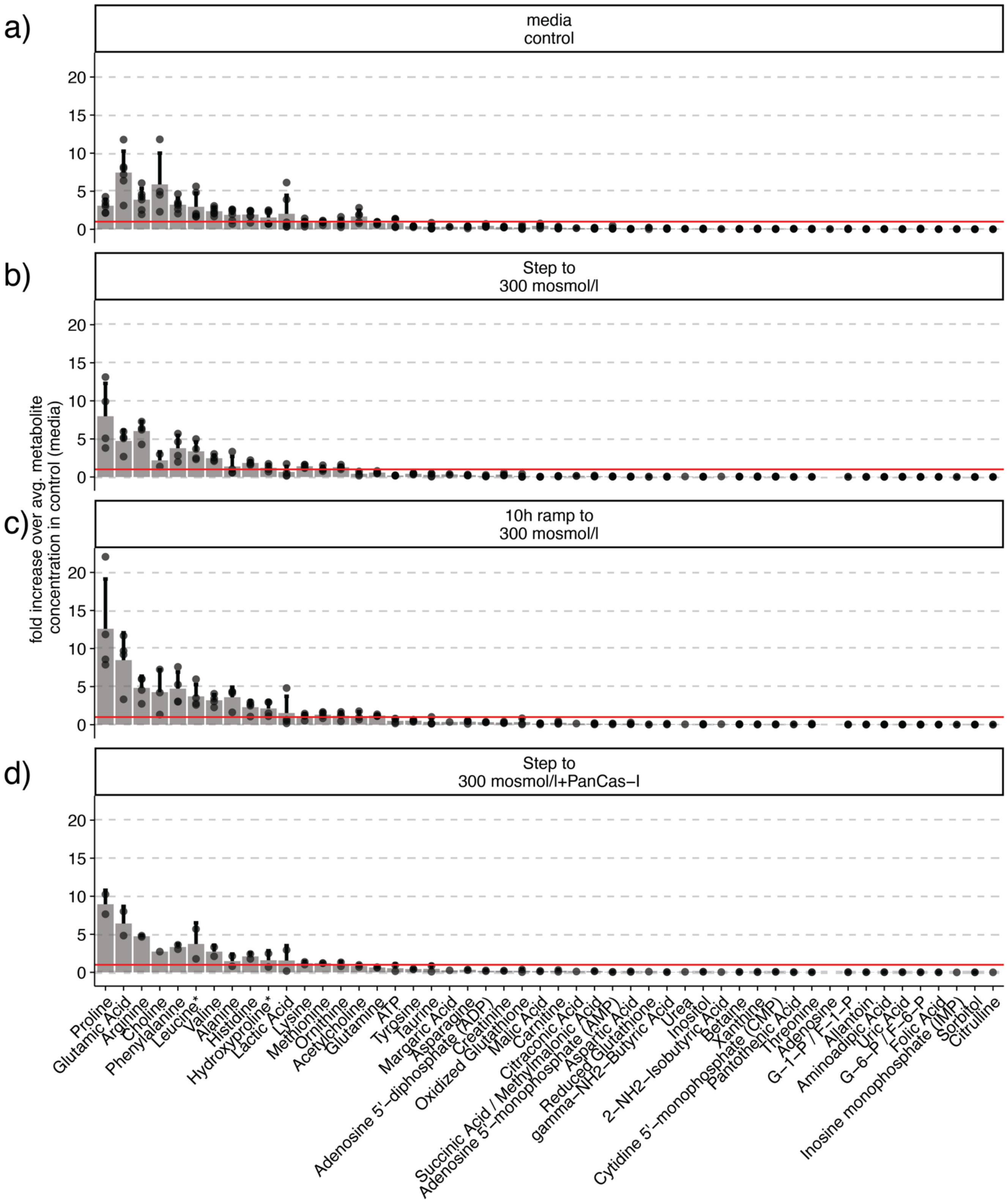


**Supplementary Figure 7: Metabolite distribution changes in hypertonic stress.** a-d) All metabolites detected in Jurkat cells (a) without stimulation, (b) after treatment with 300 mosmol/l NaCl as a step, (c) a 10h ramp, and (d) step preincubated with Pan-Caspase inhibitor (PanCas-I = 100 μM Z-VAD-FMK). Bars represent the mean and standard deviation of the fold change of each metabolite to the average metabolite concentration in the control condition. The Red line is a visual aid to identify metabolites that are above and below the average concentration in the control condition and indicates the relative average amino acid concentration.

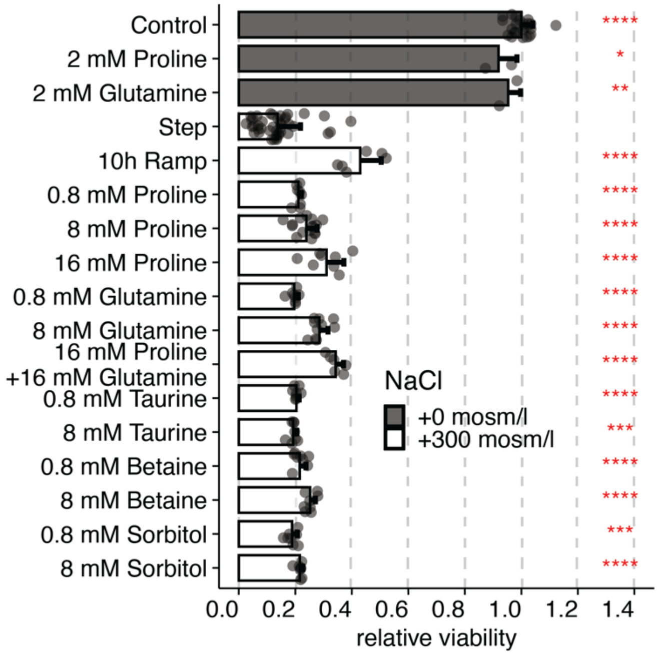


**Supplementary Figure 8: External proline and glutamine improve viability in step treated cells similarly to ramp treated cells without external proline or glutamine in comparison to established osmolytes.** Viability relative to untreated cells (control) in Jurkat cells exposed to additional 0 or 300 mosmol/l NaCl for 5h (step) or 10h (10h ramp), respectively. Bars indicate mean and SD of at least 3 replicates. Osmolytes were added 60 min before NaCl at indicated concentrations. t-test: *p<0.05, **p<0.01, ***p<0.001, ****p<0.0001.
